# RhizoGrid: Illuminating Spatial Rhizosphere Dynamics

**DOI:** 10.64898/2025.12.10.693536

**Authors:** Pubudu P. Handakumbura, Albert Rivas Ubach, Anil Battu, Winston E Anthony, Kate Schultz, Ryan McClure, Tamas Varga, Christer Jansson, Rob Egbert

## Abstract

The rhizosphere microbiome directly influences plant health and acclimation to extreme environments, yet plant-microbe interactions in the rhizosphere have proven complex and difficult to study. We present RhizoGrid, a new methodological framework that integrates a 3D-printed pot structure with spatial measurements of root structure, metabollite and taxonomy to detect links between metabolites and microbes along a soil-grown root system. The RhizoGrid identifies microhabitats hidden belowground. Using the food, forage, and bioenergy crop sorghum, we showcase how the RhizoGrid opens new frontiers for exploring the heterogeneity and complexity of interactions between root exudates and microbes in the rhizosphere.

## Main

As sessile organisms, plants are remarkably adapted to their environments and can regulate their responses to either acclimate to or escape extreme environmental changes. The soil associated with and influenced by plant roots, known as the rhizosphere, is one of the most dynamically regulated soil environments. Studies indicate that around ∼27% of the photosynthetically fixed carbon allocated to roots is released to the rhizosphere as root exudates (rhizodeposition)[1, 2] The overall composition of rhizodeposition, containing thousands of metabolites, is species-specific and, together with soil properties[3] modulate rhizosphere microbiome dynamics.[4-8] Root exudates have multiple functions, such as warning signals to prevent herbivory [9], promote symbiosis with rhizobia and mycorrhizal fungi[10], and promote beneficial microorganisms.[11, 12] Still, determining the underlying chemical signals involved in recruiting and maintaining plant species-specific root microbiomes remains a grand challenge in rhizosphere biology. [12]

The driving hypothesis for RhizoGrid development is that the plant root system produces metabolite abundance gradients,[13] creating unique, spatially discrete community assemblages via root exudate metabolites. Manipulating microbial functions in the rhizosphere to benefit plants requires a fundamental understanding of the spatiotemporal assembly of microbiomes along the root system. Most approaches homogenize the whole plant root for analysis [14], preventing assessment of the microscale spatial organization of the rhizosphere and failing to capture the unique microhabitats occupied by distinct microbial communities. The spatial impacts of metabolite rhizodeposition on bacterial community assembly is poorly understood [12, 15] and the implications for spatial enrichment of microbial community members remains an unaddressed question in rhizosphere biology.

A key challenge with the existing molecular imaging technologies at the rhizosphere scale is the lack of spatial resolution. The RhizoGrid combines cm-scale molecular level measurements from defined angular and depth grid positions to identify microenvironmental effects within the system. We have developed and optimized a root cartography workflow that leverages multidimensional multi-omics data derived from root-rhizospheres of plants grown in RhizoGrid pots to image the root system in 3-dimensional space and link spatial metabolite and microbial profiles of the rhizosphere (**Fig. 1, Fig. S1)**.[14] RhizoGrid platform consists of 3D printed pots fitted with 3D printed modular grids that can be customized for root systems of different geometries. These 3D-printed grids support the developing root system, ensuring roots grow unhindered. This approach preserves the three-dimensional architecture of a plant root system, emulating root systems observed in a field setting. Spatial indexing is built and optimized to include non-invasive imaging of the root system in the 3D space using X-ray computed tomography (XCT) coupled with harvesting and processing of the roots and the associated rhizosphere for metabolite and microbial profiling. Each excised root segment is assigned a distinct coordinate based on its RhizoGrid position. RhizoGrid-guided indexing facilitates the projection of spatially resolved omics data onto reconstructed 3D root images obtained from XCT data.

**Figure 1.**
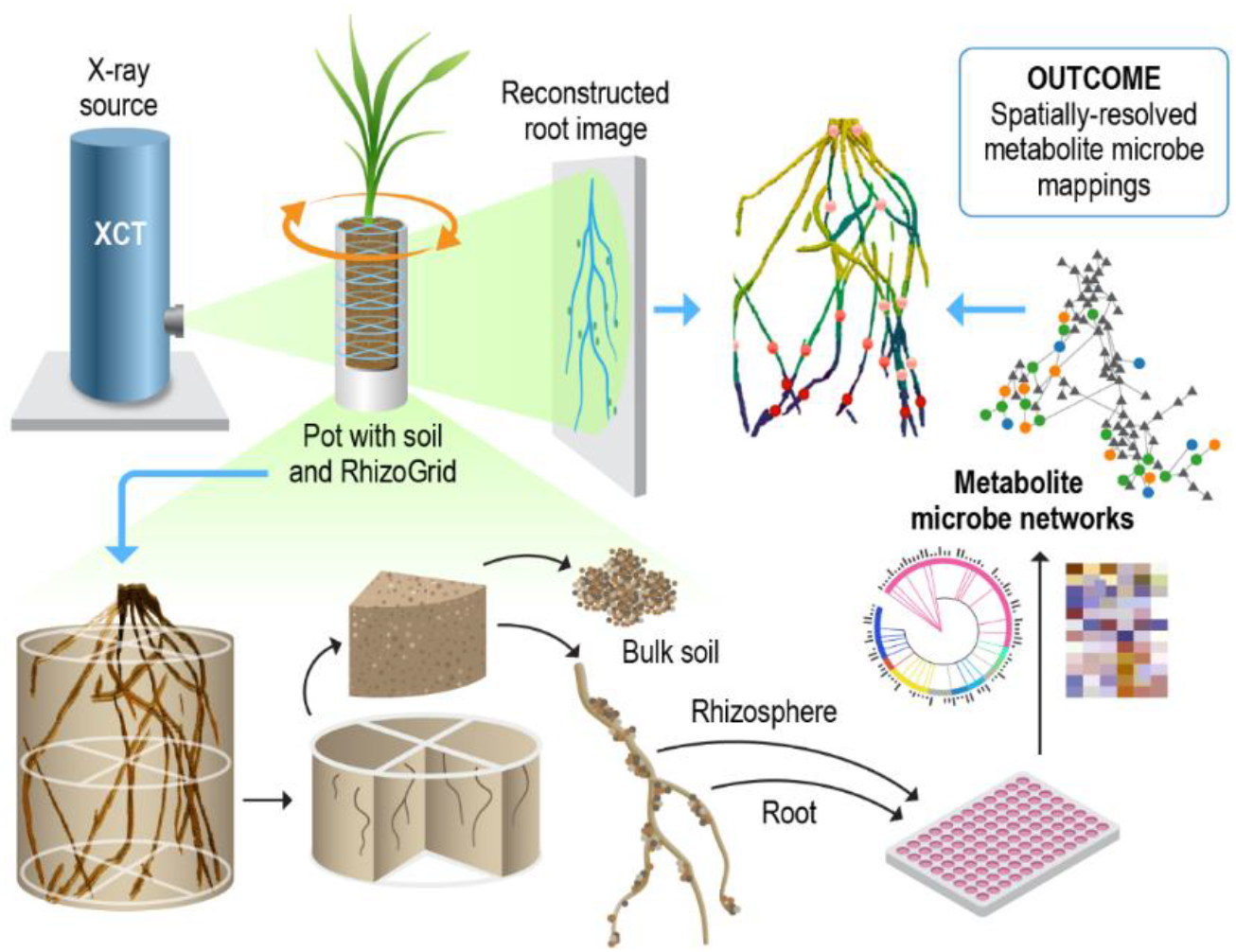
Three-dimensional root-rhizosphere cartography. A plant is grown in soil using RhizoGrid-augmented pots. A 3D image of the root is reconstructed using the image scans generated by X-ray computed tomography (XCT). Following 3D rendering of the root structure, the root and adherent soil are extracted from the pot. Loosely adhered soil is removed from the root to retain the rhizosphere. The root with its rhizosphere is sequentially harvested and separated into root and rhizosphere (root wash-off) fractions for metabolite and microbial taxonomy profiling using liquid chromatography-mass spectrometry and 16S rRNA sequencing, respectively. Metabolite and taxonomy data are mapped back to the XCT root image to visualize the rhizosphere metabolite microbe interactions.

The spatial organization of the rhizosphere is crucial to control plants’ access to nutrients, interactions with beneficial microbes, and mitigation of stress. These factors drive plant performance and ecosystem health. By creating a structured environment where different microbial populations occupy distinct niches near the root, the plant allows optimized nutrient uptake and pathogen suppression, depending on the spatial arrangement of those microbes. We utilized a local field soil from Prosser, WA, USA and the native microbiome of the soil as a microbial population to evaluate the RhizoGrid system. Sorghum seeds were directly sown into RhizoGrid-fitted pots and grown for five weeks under controlled environmental conditions in a reach-in growth chamber. Metabolites and microbial DNA was isolated from spatially sampled roots and rhizosphere of three sorghum plants following the RhizoGrid workflow. From a total of 128 16S rRNA samples, 121 sequenced samples met quality measures for retention and analysis. We recovered 5212 ASVs covering 43 phyla, 497 genera, and 140 species. 211 ASVs were present in at least half of the study samples (**Table S1**). We detected 8919 metabolite features by liquid chromatography coupled mass spectrometry (LC-MS) analyses, from which 173 metabolites were identified and a further 495 were classified to a canopus class. Among the most abundant metabolites, 49 of the top 900 (roughly 10% of 8919) were identified (**Table S2**). The top 5 identified metabolites were sucrose, malate, proline, quinate, and aconitic acid. As a result of the high complexity of the three-dimensional root system, the metabolome exhibited heterogeneity within different quadrants within the same rooting depth within a soil column, warranting further investigation.

To gain insights into the spatial arrangement of bacteria and metabolites along rooting depth, we used principal component analysis (PCA) to identify bacteria and metabolites which co-vary across depth. Meta-omic redundancy analysis (rda) of metabolomics and taxonomic measurements revealed bacteria-metabolite community correlation in 3 distinct interaction zones (**Fig. 2**). Thirteen bacterial genera and eight metabolites varied significantly by rooting depths. Bacterial abundances from the Sphingomonas, Gemmatimonas, Acidibacter, Altererythrobacter, and RB41 genera associated with the upper interaction zone. In addition, this zone was associated with increased abundances of the metabolites thymine and quinate. The middle interaction zones (representing depths 3-6 and half of depth 7) clustered together, and were associated with the microbial genera Pedomicrobium, Phyllobacterium, Nonomuraea, Allostreptomyces, and Baccillus, but no significant associations with metabolite abundances. Finally, the bottom interaction zone (lowest depth, depth 8) clustered away from the rest of the samples, and was associated with no bacterial genus, but instead with multiple metabolites, such as deoxyadenosine, umbelliferone, L hisitidine, and propanoate.

**Figure 2.**
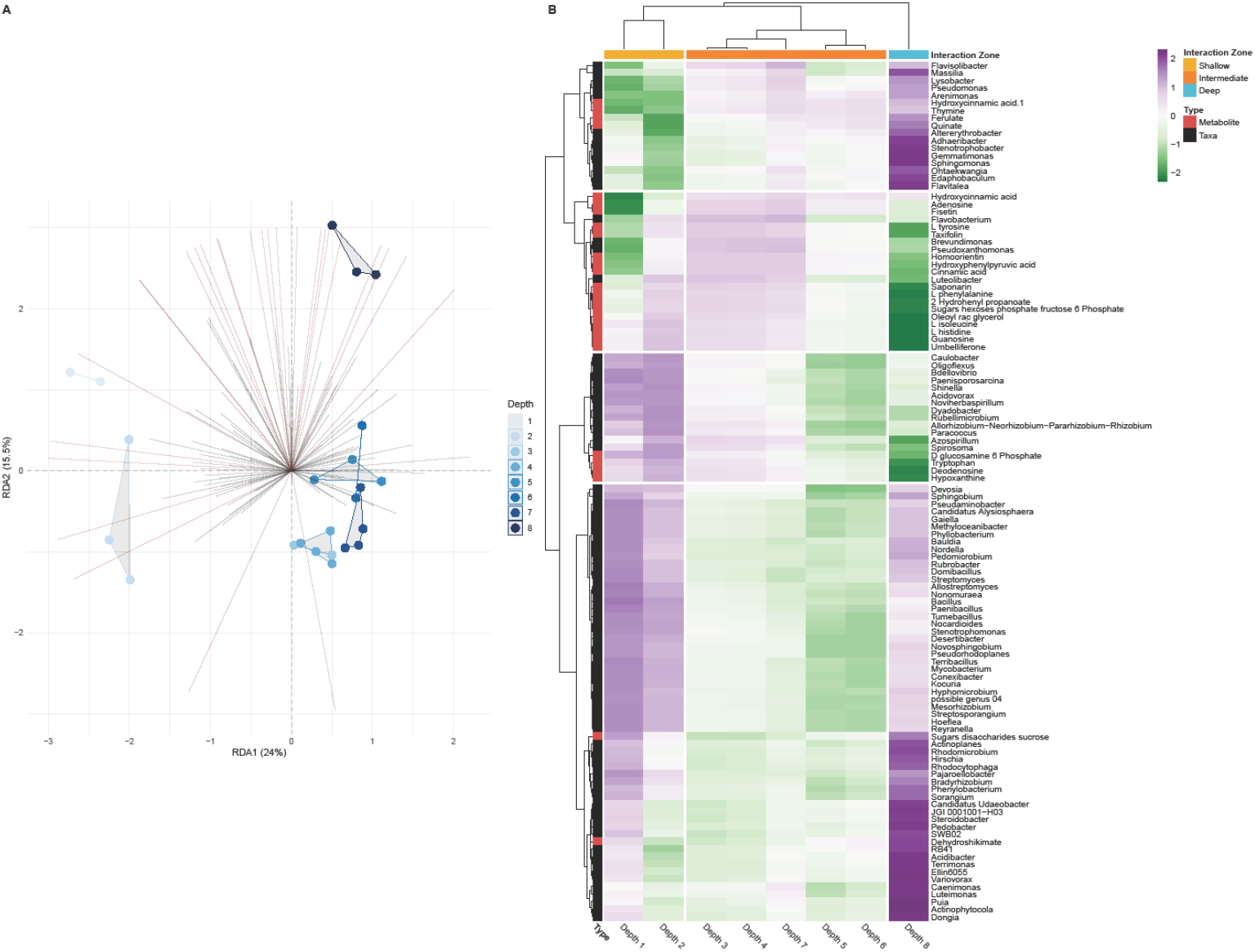
Meta-omic analysis of the rhizosphere identifies bacterial-metabolite community correlation in three separate interaction zones. A. Principal Component Analysis of 16S bacterial counts; dots represent samples colored and connected by depth. Vector loadings of significantly covarying genera and metabolites are colored black and red respectively. Samples aggregate by depth into three interaction zones. B. Bacteria and metabolite loading distance to the centroid of the samples from each depth was calculated and clustered. This resulted in 4 distinct communities which covaried across the three interaction zones: depths 1 and 2 (Shallow), depths 3-7 (Intermediate), and depth 8 (deep). Metabolite and microbial data generated from a single plant was used for this analysis.

While universal rhizosphere relationships between metabolite and microbe abundance remain complex, RhizoGrid analysis identifies metabolite-microbe covariation along depth within a single plant. The metabolites thymine and quinate were abundant in the highest root zone, and covaried with multiple bacterial genera (**Fig. 3**). Bulk microbiome samples with intact quinate degradation pathways have previously been identified in rhizosphere samples,[16] but spatially resolved RhizoGrid measurements allow a discrete, contextual understanding of the relationship between soil depth, microbial taxonomic distribution, and plant secretion products. Genera *Sphingomonas* and *Gemmatimonas* are both gram-negative rod-shaped aerobic bacteria which were present in the highest soil zone where the metabolite quinate was also in high abundance. Beyond its use as a carbon source, quinate represents an attractive intermediate in the production of shikimic acid via oxidation as it requires fewer steps than the classical pathway. Shikimic acid is a precursor for aromatic amino acids such as tyrosine, tryptophan and phenylalanine as well as secondary metabolites serving as crucial molecules in plant health, defense, and plant microbe interactions.[17] While Quinate degradation is not well characterized in these taxa, the identified association suggests an ability to degrade and utilize quinate. While Depth 8, the deepest root zone, did not associate strongly with any bacterial genus, it was enriched for several metabolites. Umbelliferone was enriched at Depth 8; it exhibits antibacterial properties against human and plant bacterial pathogens,[18] and we hypothesize that some metabolites produced at the edge of the root system may act as protective agents. Moreover, integrating multi-omics data with 3D imaging enables researchers to simultaneously analyze multiple molecular components, providing a holistic view of the root influenced rhizosphere. This comprehensive approach not only enhances our understanding of fundamental biological processes but also facilitates the identification of molecular markers associated with specific root traits. These markers are invaluable for breeding programs aimed at developing crops with enhanced traits like drought tolerance or nutrient efficiency.

**Figure 3.**
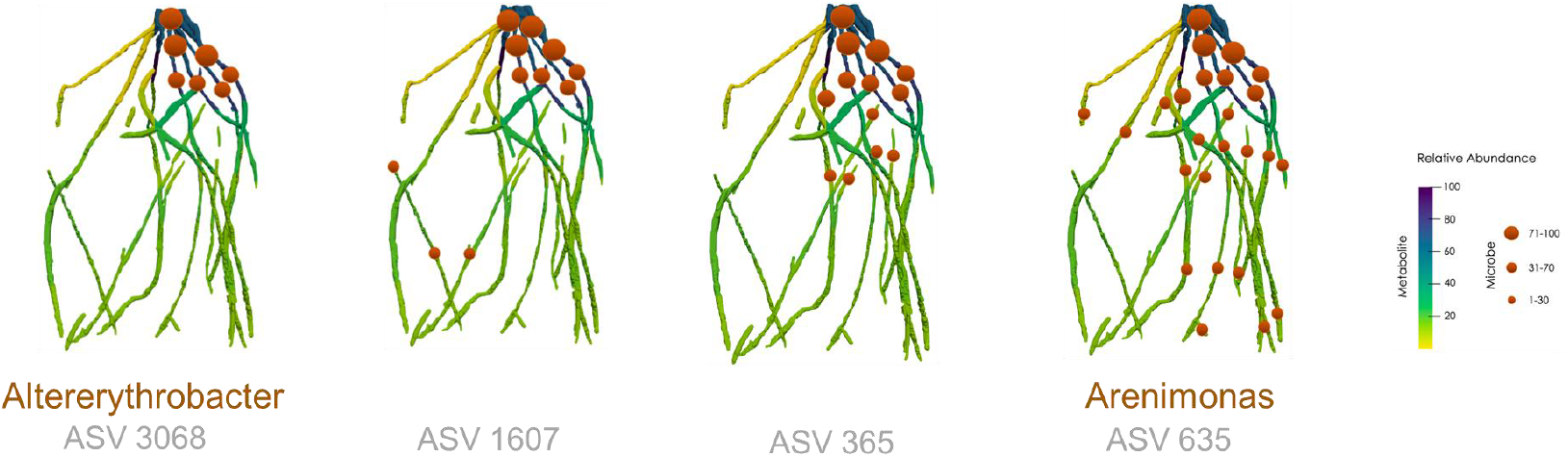
3D visualizations of selected metabolite-microbe abundances along the sorghum root system. Raw segmented images of the root and soil system were converted to 3D mesh objects and relative abundance of quinate was mapped to each segment to visualize the metabolite intensity using yellow-purple color gradient along the reconstructed Sorghum root system. Four microbial genera with significant Person correlation values were mapped to the sorghum root locations based on abundance using a red color dots with a size gradient.

Combining 3D imaging of plant roots with multi-omics data provides the research community with a powerful opportunity to explore the intricacies of root biology. However integration of 3D imaging with multi-omics data presents several challenges. These include the complexity of integrating diverse datasets from distinct experimental techniques, the difficulty of achieving high spatial resolution in both imaging and omics technologies, and the need for sophisticated computational tools and bioinformatics approaches to effectively analyze complex datasets. By correlating molecular information with specific regions of the root system, we can gain a deeper understanding of how gene expression, metabolite levels, and protein activities vary spatially. Additionally, we can leverage host-specific synthetic microbial communities to further constrain the microbial community and understand the interactions. This spatial context is crucial for unraveling complex processes such as root development, nutrient uptake, and responses to environmental stimuli. By understanding the fundamental principles governing spatial recruitment of beneficial microbiomes in the root-associated rhizosphere we can engineer synthetic rhizosphere microbiomes to sustainably promote highly productive and stress-tolerant biomass cropping systems. The RhizoGrid, an integrated imaging, and spatial multi-omics testbed, maps multi-dimensional metabolomic and taxonomic measurements to root structures. By integrating molecular and taxonomic information with 3D root images from plants grown in soil, RhizoGrid enables unprecedented access to investigate the heterogeneity and complexity of root exudate-microbial interactions in the rhizosphere.

## Author contributions

PH and CJ conceptualized the RhizoGrid workflow. PH conceived and designed the study, and wrote the original manuscript draft. PH, ARU, AB, and TV conducted the experiments. PH, ARU, RM, AB, WA, KS, and TV analyzed data, interpreted results, and revised the manuscript. RE reviewed and revised the manuscript and provided financial support.

## Acknowledgments

This research was supported by the U.S. Department of Energy (DOE), Office of Biological and Environmental Research (BER), as part of BER’s Genomic Science Program (GSP) and is a contribution of the Pacific Northwest National Laboratory (PNNL) Secure Biosystems Design Science Focus Area “Persistence Control of Engineered Functions in Complex Soil Microbiomes” and the EMSL strategic Science Area under the project 22142. A portion of this work was performed in the William R. Wiley Environmental Molecular Sciences Laboratory (EMSL), a national scientific user facility sponsored by BER and located at PNNL. PNNL is a multi-program national laboratory operated by Battelle for the DOE under Contract DE-AC05-76RL01830.

## Methods

### 1. Plant growth setup and sample collection

BTx642 sorghum seeds were directly sown into the RhizoGrid platform pots and grown for five weeks under controlled environmental conditions in a Percival reach-in growth chamber using field soil collected from a local field site in Prossor, WA. Prosser soil is characterized as a Warden fine sandy/silt loam with 0.4% organic matter and 1 mg/kg Ammonium N per dry weight. Plants were grown at 16: 8h light: dark cycles with 22: 18 °C day: night temperatures, 60 % RH, and ∼250 µmoles light intensity. The soil column was harvested sequentially from bottom to top and separated into the corresponding quadrants. Root pieces were separated from the bulk soil and rinsed in 500 ul of sterilized water and flash-frozen in LN2. Washed off soil was saved as the rhizosphere fraction.

### 2. XCT and image processing

RhizoGrid grown Sorghum plants were imaged using X-ray Computed Tomography (XCT) on an X-Tek/Metris XTH 320/225 kV scanner (Nikon Metrology, Belmont, CA). Images were collected at 100 kV and 450 μA X-ray power. A 0.25-mm thick Cu filter was used to enhance image contrast by blocking out low-energy x-rays. The samples were rotated continuously during the scans with momentary stops to collect each projection (shuttling mode) while minimizing ring artifacts. A total of 3142 projections were collected over 360° rotation recording 2 frames per projection with 708 ms exposure time per frame. Image voxel size was 41.3 microns. The images were reconstructed to obtain three-dimensional datasets using CT Pro 3D (Metris XT 2.2, Nikon Metrology). Image processing and segmentation was carried out using Avizo 2019.2 (Thermo Fisher Scientific, Waltham, MA). The reconstructed 3D volume data was filtered using a median filter, and the data was segmented into soil, root, and grid based on manual thresholding by assigning each voxel to the component it belongs to (soil, root, or grid). Root and grid were then selected from the segmented image and visualized using selected colors. Representative images were created using Avizo.

### 3. Metabolomics data acquisition and processing

Spatially sampled root tissue was lyophilized and homogenized using a SPEX SamplePrep Geno Grinder with 3mm metal beads. Every 10 mg of root tissue was extracted with 200 ul of 80:20 methanol: water, following the published protocols.[19] Sample extracts were run in liquid chromatography high-resolution mass spectrometry (LC-MS) for untargeted metabolomics analyses. Liquid chromatography (LC) was performed with a Thermo Vanquish high pressure liquid chromatographer (HPLC) (Thermo Fisher Scientific, Waltham, Massachusetts, USA). Extracted metabolites were separated by injecting 5 uL of each sample into a Hypersil gold C18 reversed-phase column (150×2.1 mm, 3 μm particle size; Thermo Scientific, Waltham, Massachusetts, USA) which was maintained constant at 30°C. Mobile phases of the LC consisted of 0.1% formic acid in water (A) and 0.1% formic acid in acetonitrile/water (90/10) (B). The elution gradient operated at a constant flow rate of 0.3 mL/min along the chromatography. The gradient started at 90% A (10% B) and was maintained stable for 5 minutes before it ramped to 10% A (90% B) until minute 20 of the chromatography. Those conditions were held constant for 2 additional minutes and the initial conditions were linearly recovered during the subsequent 2 minutes. The column was thus washed and stabilized at the initial conditions for 11 minutes before injecting the next sample (min. 35). High-resolution mass spectrometry (HRMS) analyses were carried out on a LTQ Orbitrap Velos equipped with heated electrospray ionization (HESI) source (Thermo Fisher Scientific, Waltham, Massachusetts, USA). Each sample was run at both positive and negative ionization modes. The HRMS operated at full-scan mode, at a resolving power of 60,000 full width at half maximum (FWHM) at 400 m/z and tandem mass spectrometry (acquisition of MS1 and MS/MS data).Ions were between 70 and 800 m/z. Data dependent acquisition (DDA) was set to select the top 3 most intense precursor ions of each scan and obtain MS/MS spectra. MS/MS data was acquired in the Orbitrap at a resolving power of 7.500. LC-MS RAW files acquired in negative and positive ionization modes were processed in MZmine 2.53.[20] Each ionization mode was processed independently. Briefly, for each sample, chromatographic baseline was corrected and metabolite features with isolated m/z and RT values were detected, deconvoluted, aligned and gap filled. Based on the RT and exact mass, deconvoluted features were matched against an in-house metabolite library and peak areas corresponding to MS1 were thus exported to a CSV file. Using both RT and the exact mass of detected ions to match against metabolite libraries represent a second level of putative identification.[21] MS/MS data of each ionization mode was separately exported to a MGF file and subsequently processed in SIRIUS 4[22] here compound classes were assigned to each individual feature with MS/MS data using the CANOPUS module[23] MS1 peak areas were filtered before statistical analyses. Briefly, data present in experimental blanks was used to determine the noise level of the instrument. The noise of each signal individually was calculated averaging the peak areas of each blank for such specific feature. If the number of blanks with data in a specific feature was lower than 30%, the noise level was considered as zero. Then, for each feature and sample, only those features with a signal to noise (S/N) equal or larger than 10 were considered in the study.

### 4. Microbial DNA isolation and 16s data acquisition

DNA extraction was conducted using Quick-DNATM Fecal/Soil Microbe Miniprep Kit (D6010) and samples were prepared according to the kit protocol. Samples were cleaned and concentrated with a kit (Zymoreserach ZR-96 Genomic DNA Clean & Concentrator *https://www.zymoresearch.com/products/zr-96-genomic-dna-clean-concentrator-5-kit).* Raw fastq files were examined with Hundo to assign them to taxonomies and were normalized via rarefaction (using the rrarefy function in R). Any OTU that had a total count of less than 10 (across all samples) was removed from analysis and biological replicates were averaged. To create a network of bacterial genera and metabolites, average normalized counts for each genus were generated by averaging the normalized counts for all OTUs in the given genus. This genus-averaged data was then combined with metabolite data. Only known metabolites were used for network analysis. Pearson correlation coefficient was used to link metabolites and genera. Only genera and metabolites that had an absolute value correlation of >0.65 were used to infer a network. Magnitude of the correlation as well as the direction was then overlayed onto the network and displayed as line thickness or color, respectively.

### 5. Redundancy analysis of combined omics data

Rarified 16S genera counts for 1 plant were first subjected to a prevalence threshold where genera not found in at least 3 individual sampling sites were removed from analysis. The data was then transformed with the robust centered-log ratio transformation using the ‘rclr’ option in the vegdist function from the vegan statistical analysis package in r. RDA was performed on these data using the rda function from vegan and genus loading vectors with a coefficient of determination (r2) above 70% and p < 0.01 were plotted. Using the subset of positively identified metabolomic trace data (after data was transformed using the Hellinger transformation using the same vegan function listed above) from the same sampling sites, vector fitting was conducted and the metabolomic vectors with r2 values above 60% and p < 0.01 were plotted. Data was extracted from r objects using the BiodiversityR package and visualized using ggplot2.

### 6. Spatially resolved metabolite-microbe mappings onto the sorghum root system

Raw segmented images of the root and soil system were converted to 3D mesh objects using the Python library PyMesh (https://pymesh.readthedocs.io/). Each of the 32 quadrants used during sampling were then mapped to the root mesh, forming a 3D grid with each of the 32 segments defined according to its center point. Finally, the mesh was annotated to assign each quadrant values corresponding to relative abundance data for a given metabolite or microbial species. These values were converted to a designated color scale to color each root segment accordingly, with the resulting files loaded into visualizer software as.ply files. A Python notebook with code for creating these visualizations with PyMesh is available in the Supporting Information.

---

**Figure S1.**
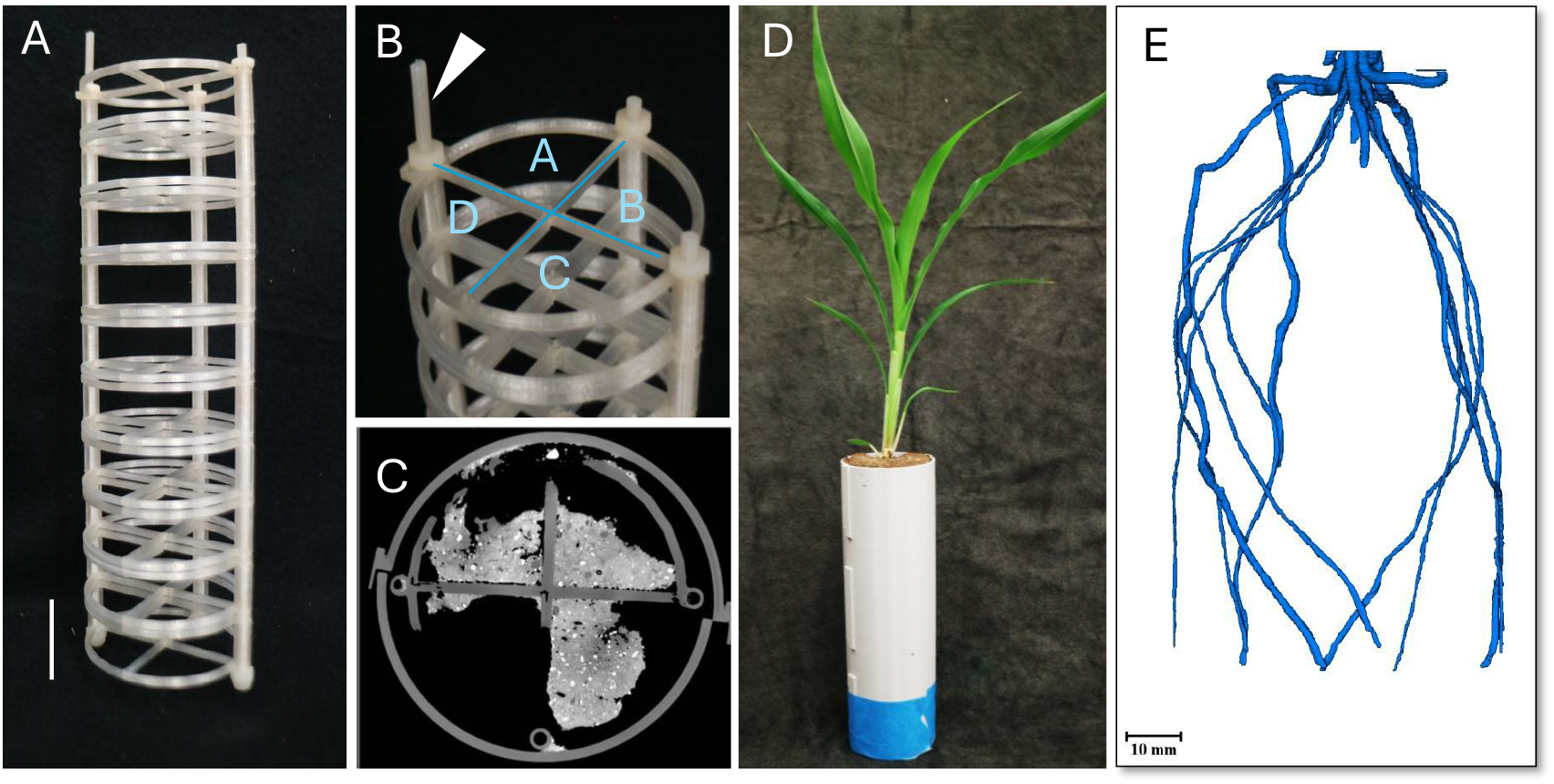
RhizoGrid Platform. **a**. 3D printed customizable grids (scale bar = 1 inch), **b**. Top view of the pot with the A-D quadrants marked for spatial localization of root samples within the three-dimensional space. White arrowhead marks the longer rod used to mark the starting position for quadrant labeling **c**. Arial view of the pot in an XCT scan illustrating the contrast between the 3D printed grids and the soil within the pot, **d**. The sorghum plant grown in field soil collected from a local field site in Prosser, Washington, **E**. 3D reconstruction of soil-grown sorghum root using XCT images (root false colored in blue).

## References

1. York LM, Carminati A, Mooney SJ, Ritz K, Bennett MJ: The holistic rhizosphere: integrating zones, processes, and semantics in the soil influenced by roots. J Exp Bot 2016, 67(12):3629–3643.

2. Jones DL, Hodge A, Kuzyakov Y: Plant and mycorrhizal regulation of rhizodeposition. New Phytol 2004, 163(3):459–480.

3. Koebernick N, Daly KR, Keyes SD, George TS, Brown LK, Raffan A, Cooper LJ, Naveed M, Bengough AG, Sinclair I et al: High-resolution synchrotron imaging shows that root hairs influence rhizosphere soil structure formation. New Phytol 2017, 216(1):124–135.

4. Massalha H, Korenblum E, Tholl D, Aharoni A: Small molecules below-ground: the role of specialized metabolites in the rhizosphere. Plant J 2017, 90(4):788–807.

5. Vorholt JA, Vogel C, Carlstrom CI, Muller DB: Establishing Causality: Opportunities of Synthetic Communities for Plant Microbiome Research. Cell Host Microbe 2017, 22(2):142–155.

6. Garbeva P, van Elsas JD, van Veen JA: Rhizosphere microbial community and its response to plant species and soil history. Plant and Soil 2008, 302(1):19–32.

7. Jiang Y, Li S, Li R, Zhang J, Liu Y, Lv L, Zhu H, Wu W, Li W: Plant cultivars imprint the rhizosphere bacterial community composition and association networks. Soil Biology and Biochemistry 2017, 109:145–155.

8. Bulgarelli D, Schlaeppi K, Spaepen S, Ver Loren van Themaat E, Schulze-Lefert P: Structure and functions of the bacterial microbiota of plants. Annu Rev Plant Biol 2013, 64:807–838.

9. Babikova Z, Gilbert L, Bruce TJ, Birkett M, Caulfield JC, Woodcock C, Pickett JA, Johnson D: Underground signals carried through common mycelial networks warn neighbouring plants of aphid attack. Ecol Lett 2013, 16(7):835–843.

10. Badri DV, Vivanco JM: Regulation and function of root exudates. Plant Cell Environ 2009, 32(6):666–681.

11. Pang Z, Chen J, Wang T, Gao C, Li Z, Guo L, Xu J, Cheng Y: Linking Plant Secondary Metabolites and Plant Microbiomes: A Review. Front Plant Sci 2021, 12:621276.

12. Zhalnina K, Louie KB, Hao Z, Mansoori N, da Rocha UN, Shi S, Cho H, Karaoz U, Loque D, Bowen BP et al: Dynamic root exudate chemistry and microbial substrate preferences drive patterns in rhizosphere microbial community assembly. Nat Microbiol 2018, 3(4):470–480.

13. Zhang T, Noll SE, Peng JT, Klair A, Tripka A, Stutzman N, Cheng C, Zare RN, Dickinson AJ: Chemical imaging reveals diverse functions of tricarboxylic acid metabolites in root growth and development. Nature Communications 2023, 14(1):2567.

14. Handakumbura PP, Rivas Ubach A, Battu AK: Visualizing the Hidden Half: Plant-Microbe Interactions in the Rhizosphere. mSystems 2021, 6(5):e0076521.

15. Jacoby RP, Kopriva S: Metabolic niches in the rhizosphere microbiome: new tools and approaches to analyse metabolic mechanisms of plant-microbe nutrient exchange. J Exp Bot 2019, 70(4):1087–1094.

16. Singh DP, Prabha R, Gupta VK, Verma MK: Metatranscriptome Analysis Deciphers Multifunctional Genes and Enzymes Linked With the Degradation of Aromatic Compounds and Pesticides in the Wheat Rhizosphere. Front Microbiol 2018, 9:1331.

17. Yokoyama R, de Oliveira MVV, Kleven B, Maeda HA: The entry reaction of the plant shikimate pathway is subjected to highly complex metabolite-mediated regulation. Plant Cell 2021, 33(3):671–696.

18. Soares V, Marini MB, de Paula LA, Gabry PS, Amaral ACF, Malafaia CA, Leal ICR: Umbelliferone esters with antibacterial activity produced by lipase-mediated biocatalytic pathway. Biotechnology Letters 2021, 43(2):469–477.

19. Handakumbura PP, Stanfill B, Rivas-Ubach A, Fortin D, Vogel JP, Jansson C: Metabotyping as a Stopover in Genome-to-Phenome Mapping. Sci Rep 2019, 9(1):1858.

20. Pluskal T, Castillo S, Villar-Briones A, Orešič M: MZmine 2: Modular framework for processing, visualizing, and analyzing mass spectrometry-based molecular profile data. BMC Bioinformatics 2010, 11(1):395.

21. Sumner LW, Amberg A, Barrett D, Beale MH, Beger R, Daykin CA, Fan TWM, Fiehn O, Goodacre R, Griffin JL et al: Proposed minimum reporting standards for chemical analysis. Metabolomics 2007, 3(3):211–221.

22. Dührkop K, Fleischauer M, Ludwig M, Aksenov AA, Melnik AV, Meusel M, Dorrestein PC, Rousu J, Böcker S: SIRIUS 4: a rapid tool for turning tandem mass spectra into metabolite structure information. Nature Methods 2019, 16(4):299–302.

23. Dührkop K, Nothias L-F, Fleischauer M, Reher R, Ludwig M, Hoffmann MA, Petras D, Gerwick WH, Rousu J, Dorrestein PC et al: Systematic classification of unknown metabolites using highresolution fragmentation mass spectra. Nature Biotechnology 2021, 39(4):462–471.

